# Cryo-EM structure of a RAS/RAF recruitment complex

**DOI:** 10.1101/2022.07.14.500081

**Authors:** Eunyoung Park, Shaun Rawson, Anna Schmoker, Byeong-Won Kim, Sehee Oh, KangKang Song, Hyesung Jeon, Michael Eck

## Abstract

Cryo-EM structures of a KRAS/BRAF/MEK1/14-3-3 complex reveal KRAS bound to the flexible Ras-binding domain of BRAF, captured in two orientations. Autoinhibitory interactions are unperturbed by binding of KRAS and *in vitro* activation studies confirm that KRAS binding is insufficient to activate BRAF, absent membrane recruitment. These structures illustrate the separability of binding and activation of BRAF by Ras and suggest stabilization of this pre-activation intermediate as an alternative to blocking binding of KRAS.

---

RAF-family kinases are a central point of control in the signaling apparatus that regulates cellular proliferation, growth and differentiation^1,2^. RAFs are maintained in an inactive, autoinhibited state in the cytosol and are activated downstream of receptor tyrosine kinases via recruitment to the plasma membrane by GTP-bound RAS. Upon activation, RAFs phosphorylate their sole known substrate MEK1/2, and activated MEK in turn phosphorylates ERK1/2. Somatic mutations in RAF can subvert the normal RAS-dependent activation process and are a frequent cause of cancer^3,4^. In particular the V600E mutation in BRAF drives more than half of malignant melanoma and is also found in lung, colorectal, thyroid, and many other cancers^5^. The conserved domain structure of BRAF and other family members (ARAF and CRAF, also called RAF-1) includes the RAS binding domain (RBD), a cysteine-rich domain (CRD), and the C-terminal kinase domain (Fig. 1a). The RBD and CRD lie in the N-terminal regulatory region of the protein, which is crucial both for maintaining autoinhibition and for RAS-driven activation and membrane association^1,2^. We and others have shown how BRAF is locked in an inactive state as a complex with its substrate MEK and a 14-3-3 dimer^6,7^. In this autoinhibited configuration, the 14-3-3 dimer binds serine phosphorylation sites that flank the BRAF kinase domain (pS365 and pS729), allowing it to restrain both the BRAF kinase and CRD domains in a cradle-like structure that precludes BRAF dimerization, which is crucial for its activation^8^. The CRD domain lies at the center of the autoinhibited complex, where it is largely protected from interactions with the membrane and RAS. The RBD, by contrast, is exposed and available to engage RAS. In the active state, driven by Ras-engagement at the plasma membrane, these components reorganize such that the 14-3-3 domain binds the C-terminal pS729 site of two BRAF proteins, driving formation of an active, back-to-back BRAF kinase dimer^6,9,10^. The N-terminal regulatory region appears not to engage with the dimeric kinase/14-3-3 module^6,9^; instead, it is thought to bind RAS at the membrane^11-13^.

**Figure 1.**
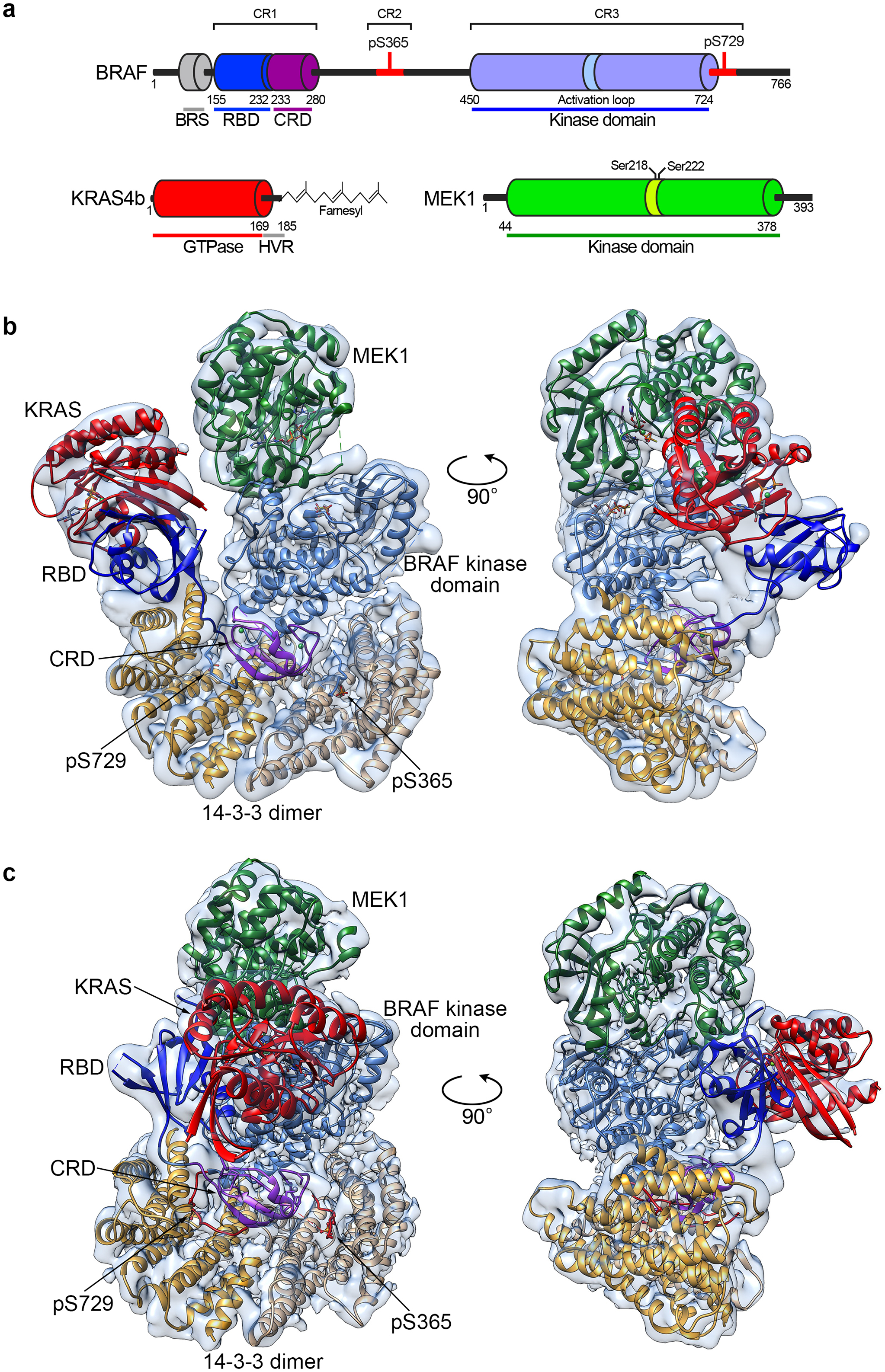
Cryo-EM structures of a KRAS/BRAF/MEK1/14-3-3 complex. a, Schematic of domain structures of BRAF, KRAS, and MEK1. Key phosphorylation sites are indicated above the schematics, and residue numbers for domain boundaries are shown below. Binding sites for the 14-3-3 domain in BRAF are indicated in red. Abbreviations: BRS, BRAF-specific domain; RBD, Ras binding domain; CRD, cysteine-rich domain; HVR, Ras hypervariable region. b, Structure of the KRAS/BRAF/MEK1/14-3-3 complex in the “Ras up” conformation. A ribbon diagram of the structure, colored as in panel a above, is shown together with the transparent cryo-EM density map. c, Structure of the KRAS/BRAF/MEK1/14-3-3 complex in the “Ras front” conformation, determined with the BS3 crosslinked sample. A ribbon diagram of the structure, colored as in panel a above, is shown together with the transparent cryo-EM density map.

Structural studies of RAS bound to RAFs have focused primarily on the isolated RAF RBD domain^14,15^, and more recently on a fragment of RAF-1 containing both the RBD and CRD domains^16-18^. These studies have provided a detailed view of intermolecular interactions between these domains, along with insights into how they may assemble in a membrane context^16^. In brief, RAS proteins engage the RBD domain largely via their switch I region, including an extensive β-strand interaction with the β2-strand of the RBD. RAS also engages the adjacent CRD domain, although the interaction apparently contributes little to binding affinity. The CRD domain therefore has a dual role in the active state, in that it binds both the membrane and RAS. Despite this detailed understanding of KRAS/RAF recognition, it remains unclear how this interaction results in RAF activation. As a next step in our effort to structurally dissect this process, we have determined the structure of KRAS bound to the full-length BRAF/MEK/14-3-3 complex using cryo EM. The structures reveal a pre-activation intermediate, in which key autoinhibitory interactions are unperturbed, despite engagement of KRAS.

Addition of the KRAS GTPase domain (KRAS^GTPase^, residues 1-169, stabilized in its active state by loading with the non-hydrolyzable GTP analog GMP-PNP), to the autoinhibited BRAF/MEK/14-3-3 complex resulted in formation of a stable complex, as judged by co-elution on size-exclusion chromatography (Extended Data Fig. 1). The elution volume was little changed as compared with that of the autoinhibited BRAF complex alone, suggesting that addition of KRAS had not triggered conversion to the active dimeric state. A single particle cryo EM reconstruction at ∼4.3Å resolution revealed that the complex indeed retains an autoinhibited configuration, with KRAS bound to the BRAF RBD domain and extending “up” from the 14-3-3 dimer, positioning KRAS alongside the C-lobe of the MEK1 kinase domain (Fig. 1b, Extended Data Table 1). The density for both KRAS and the BRAF RBD was somewhat weaker and lower resolution than the remainder of the complex (Extended Data Fig. 2a), indicative of conformational variability of these domains, but we were nevertheless able to unambiguously position both domains (Fig. 1b). The interaction between KRAS and the BRAF RBD in this structure is analogous that previously seen in crystal structures of CRAF bound to RAS^15,17^; KRAS uses primarily its “switch I” region to bind the RBD, forming an anti-parallel β-strand interaction with the β2-strand of the RBD domain (Extended Data Fig. 3a,b). KRAS makes only a glancing contact with the MEK; both its N- and C-termini are positioned within 6-7Å of the MEK C-lobe, near residues Gly237 to Tyr340 in the MEK1 C-lobe (Extended Data Fig. 3c).

We also imaged a similar sample that was subjected to crosslinking with Bis(sulfosuccinimidyl)suberate (BS3) prior to size-exclusion chromatography. Using three-dimensional classification, we obtained two reconstructions with this sample, one in a “RAS-up” conformation as described above, and the other in a “RAS-front” conformation in which the KRAS/RBD module pivots to position KRAS in front of the BRAF and MEK1 kinase domains (Fig. 1c, Extended Data Table 1, Extended Data Fig. 2b). The RAS-front conformation is not induced by crosslinking, as we also observed this conformation with three-dimensional classification of particles in the non-crosslinked sample (Extended Data Fig. 2a). Overall, the resolution of the cross-linked RAS-front reconstruction is ∼3.9Å, but as in the non-crosslinked structure, the resolution in the region of the KRAS/RBD is considerably lower. In this conformation, KRAS binds the RBD in the general manner expected, but there is approximately a 11° difference in their relative orientation (Extended Data Fig. 3d). Both the RBD and KRAS contact the interface between the C-lobes of the BRAF and MEK kinase domains (Fig. 1c, Extended Data Fig. 3e), but given the limited resolution in this region it is not possible to define specific interactions. Indeed, considering the apparent flexibility of the KRAS/RBD unit in both structures, we do not ascribe a particular significance to either the Ras-up or Ras-front conformations. Rather, they appear to be two orientations of the mobile KRAS/RBD module that are sufficiently populated to allow 3D-reconstruction.

We further probed inter-domain interactions in this complex with mass spectrometry analysis of the BS3-crosslinked sample (Supplementary Data File 1, Extended Data Fig. 4). We identified only one inter-domain crosslink to KRAS (connecting Lys147 in KRAS with Lys353 in MEK1) and one with the BRAF RBD domain (connecting Lys 183 in the RBD to Lys 253 in the CRD domain). These crosslinks are inconsistent with both the RAS-up and RAS-front conformations in our cryo EM reconstructions (Extended Data Fig. 4a,b), and thus are suggestive of flexibility or additional positions of the KRAS/RBD region. We do observe two crosslinks just N-terminal to the RBD domain that are consistent with the RAS-front conformation (connecting Lys150 in BRAF to MEK residues Lys185 and Lys 353).

While the RBD was exposed and poorly ordered in our prior cryo-EM structure of autoinhibited BRAF^6^, recent structures autoinhibited BRAF from Martinez Fiesco et al. reveal the RBD in a defined orientation in which binding of KRAS would lead to modest steric clashes with the 14-3-3 domain^7^. Furthermore, crystal structures of the CRAF RBD/CRD region in complex with KRAS and HRAS show essentially identical interactions of the CRD domain with RAS in addition to the well-characterized primary interaction with the RBD^17,18^. To better understand determinants of KRAS binding, we measured its affinity for various fragments of BRAF and for full-length BRAF in both autoinhibited and active states using microscale thermophoresis (Extended Data Fig. 5). The affinity of GMP-PNP loaded KRAS for full length BRAF in the active, dimeric state (∼106 nM) was similar to that in the autoinhibited state (∼126 nM), and essentially identical to its affinity for an isolated N-terminal fragment containing both the RBD and CRD domains (∼108 nM). For the isolated RBD domain, we measured an affinity of ∼21 nM. These measurements are in good agreement with previous studies of H-Ras and KRAS binding to various BRAF constructs^7,17,19^ and they point to the RBD domain as the primary energetic driver of BRAF binding to KRAS. Furthermore, they indicate that the position and interactions of the RBD in the autoinhibited state introduce little if any energetic barrier to binding of KRAS.

KRAS and other Ras isoforms are modified at their C-terminus by sequential attachment of a farnesyl or geranylgeranyl group, cleavage of the three C-terminal residues, and carboxymethylation^20^. Prenylation promotes membrane-association of Ras, and has long been known to be a requirement for its activation of Raf ^21,22^. Thus it is not surprising that the RAF/MEK/14-3-3 complex is maintained in its autoinhibited configuration when bound to KRAS^GTPase^, as this KRAS construct lacks the C-terminal hypervariable region and prenylation site. To further explore Raf activation by KRAS *in vitro*, we developed a reconstitution assay in which activation of the autoinhibited BRAF/MEK/14-3-3 complex is detected by measurement of Erk1 phosphorylation. This assay employs autoinhibited complexes prepared with MEK1^WT^, rather than the MEK1^SASA^ mutant, and thus does not require exogenous MEK1. In addition to GMP-PNP KRAS^GTPase^, we prepared intact KRAS in its fully modified, farnesylated state (KRAS4b-FME)^23^. As expected, we did not observe detectable activation by GMP-PNP KRAS^GTPase^ in this assay (Fig. 2a). Addition of KRAS4b-FME induced only a very modest and somewhat variable degree of activation (Fig. 2a,b). Addition of phosphoserine-containing liposomes to the autoinhibited RAF/MEK/14-3-3 complex also resulted in a modest increase in Erk1 phosphorylation, and the combination of liposomes and KRAS4b-FME yielded more robust activation (Fig. 2b). However, the level of activity observed with liposome-associated KRAS4b-FME was markedly less than that of purified active, dimeric BRAF (Fig. 2c).

**Figure 2.**
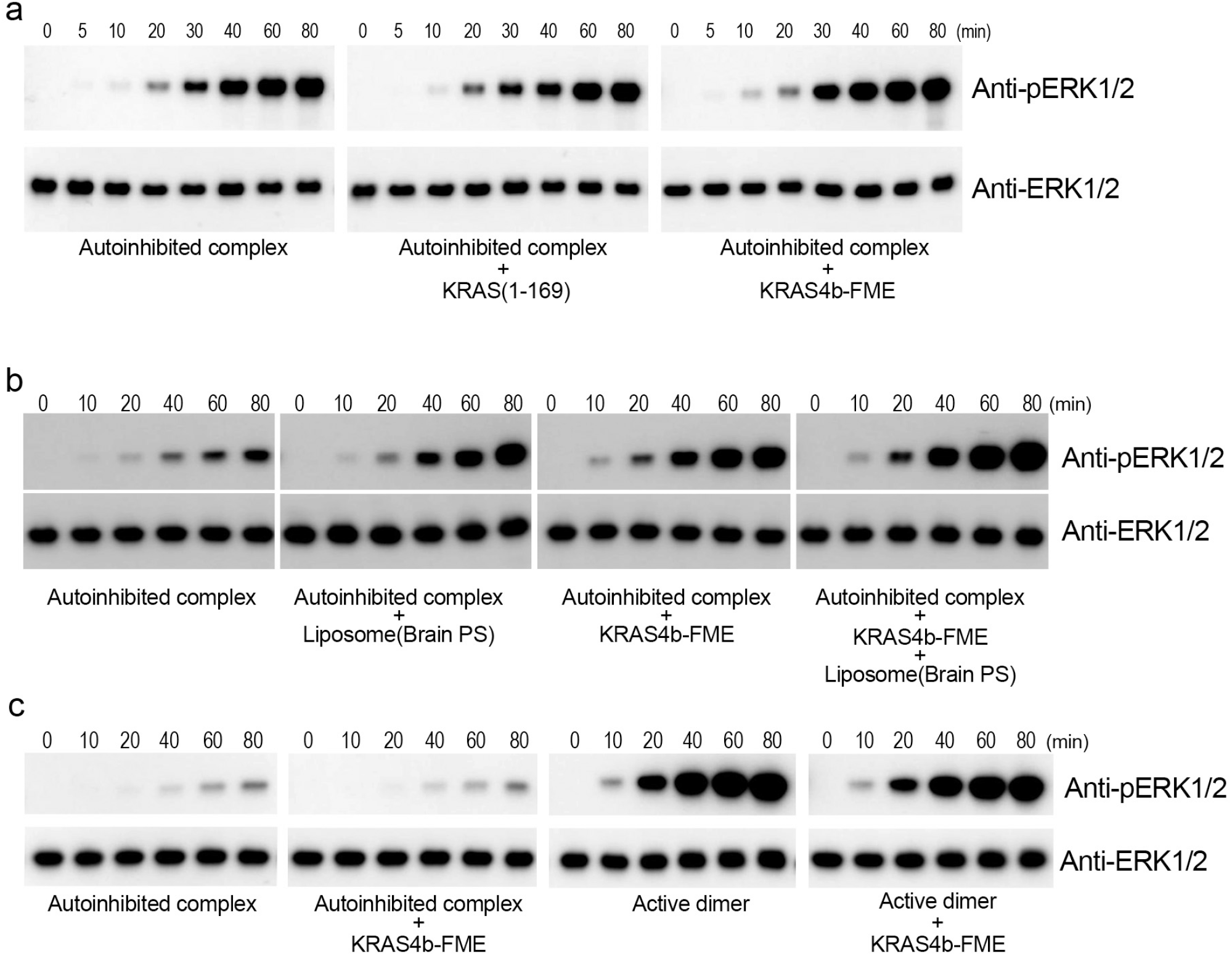
Activation studies of the autoinhibited BRAF/MEK1/14-3-3 complex. In a-c, the time course of phosphorylation of Erk1 by the purified BRAF/MEK1/14-3-3 complex, with or without addition of the indicated KRAS construct and/or liposomes, is measured by western blotting with an anti-pERK1/2 antibody. a, Activity of the autoinhibited complex alone, or with addition of either GMP-PNP loaded KRAS^GTPase^ or KRAS4b-FME at a 1:1 molar ratio. b, Activity of the autoinhibited complex alone or with addition of phosphoserine (PS) liposomes, KRAS4b-FME, or both liposomes and KRAS4b-FME, as indicated. Liposomes were added to a final concentration of 0.1 mg/ml and GMP-PNP loaded KRAS4b-FME was added in a 2:1 molar ratio to the autoinhibited BRAF/MEK1/14-3-3 complex (5 nM). c, Comparison of the relative activity of the BRAF/MEK1/14-3-3 complex in the autoinhibited vs. active, dimeric state, with or without addition of KRAS4b-FME. The samples for the autoinhibited complex with and without KRAS4b-FME are aliquots of the same reaction as the corresponding samples in panel b, re-run on the same gel and blot with the active dimer reactions to allow comparison of their relative activities. Note that the exposure of the blot in panel c (5 seconds) is much shorter than that in panel b (2 minutes).

Raf activation is a complex, multistep process. Considering that the KRAS-engaged BRAF complex maintains its autoinhibited configuration, the structures described here are best considered as “recruitment complexes” or views of a pre-activation intermediate in this process. While we observe some activation of the autoinhibited complex in *in vitro* reconstitutions with KRAS-FME in a membrane environment, activation appears to be incomplete, as judged by activity of purified BRAF/14-3-3 dimers^6^. These reconstitutions provide a starting point for incorporation of additional factors that contribute to activation. In particular, the SHOC2 phosphatase complex is thought to participate in Raf activation via dephosphorylation of the autoinhibitory 14-3-3 binding site (pS365 in BRAF, pS259 in CRAF) ^24,25^. Kinase suppressor of Ras proteins (KSR1/2) may also contribute to BRAF activation via heterodimerization and interactions with the N-terminal BRAF-specific domain^1,26^.

The structures described here clearly show that engagement of BRAF by Ras does not directly disrupt the autoinhibited complex. Thus binding and activation by KRAS are distinct steps. Considerable effort has been devoted to blocking engagement of Raf and other effectors by RAS, with little success apart from recently developed covalent KRAS G12C inhibitors^27^. This is not surprising, as high affinity protein-protein interactions are notoriously difficult to disrupt with small molecule inhibitors. The separability of KRAS binding and RAF activation illustrated here suggests stabilization of the recruitment complex as a potentially attractive alternative approach. A small molecule with a “splinting” or “molecular glue” mechanism ^28^ that stabilized interactions of the 14-3-3 domain with the kinase C-lobe or CRD domain could block RAF activation by preventing opening of the autoinhibited state, which is a prerequisite for pS365 dephosphorylation and RAF dimerization and activation.

## Acknowledgements

We thank the staff of the Harvard Cryo-EM Center for Structural Biology for assistance with data collection. This work was supported by National Institutes of Health grants R35CA242461 (M.J.E.), PO1CA154303 (M.J.E.), P50CA165962 (M.J.E.) and R50CA221830 (E.P.), and by the PLGA fund of the Pediatric Brain Tumor Foundation.

## Data Availability

Cryo-EM density maps for the structures described here have been deposited in the Electron Microscopy Data Bank (EMDB) and are available at https://www.emdataresource.org/ with accession codes EMD-27428 (KRAS-up structure) and EMD-27429 (KRAS-front structure). The corresponding atomic coordinates have been deposited in the Protein Data Bank and are available at www.rcsb.org with accession codes 8DGS and 8DGT.

## Author Contributions

E.P. expressed and purified the KRAS and BRAF/MEK/14-3-3 complexes and designed and executed the crosslinking and KRAS/BRAF/MEK/ERK pathway activation studies described here, with assistance from S.O. S.R. carried out the single particle reconstructions. H.J. prepared cryo-EM samples and collected EM data; K.S. collected cryo-EM data. A.S. executed and interpreted the crosslinking mass spectrometry study. B-W.K. designed and executed the microscale thermophoresis binding studies. E.P. and M.J.E. built the atomic models. H.J. and M.J.E. directed the project, and M.J.E. drafted the manuscript with input from all authors.

## Methods

### Preparation of KRAS proteins

Recombinant KRAS^GTPase^ (residue 1-169) was prepared as described previously^17^. For production of full-length farnesylated and methylated KRAS4b (KRAS-FME), a customized baculoviral expression system was obtained from D. Esposito (NCI-Frederick), allowing isolation of the fully processed KRAS4b protein from infected Sf9 insect cells as described previously^23^.

### Preparation of KRAS^1-169^/BRAF^WT^/MEK1^SASA^/14-3-3 complex

The BRAF^WT^/MEK1^SASA^/14-3-3 complex was prepared in an autoinhibited state as described^6^. GMP-PNP loaded KRAS^GTPase^ was added to the purified BRAF^WT^/MEK1^SASA^/14-3-3 complex in a 1.2-fold molar excess. The mixture was incubated for 30 min on ice, then applied to a size-exclusion chromatography column (Superose 6 increase 10/300, Cytiva) to remove excess KRAS protein. Size-exclusion chromatography was performed in SEC buffer (50 mM Tris pH 7.5, 150 mM NaCl, 2 mM MgCl_2_, 0.5 mM TCEP, 10 μM ATP-γS, 2 μM GDC-0623, 10 μM GMP-PNP).

### Preparation of cross-linked KRAS^GTPase^/BRAF^WT^/MEK1^SASA^/14-3-3 complex

KRAS^GTPase^ was added to the BRAF autoinhibited complex as described above, incubated on ice for 30 min, then loaded onto size-exclusion chromatography in crosslinking buffer (20 mM HEPES pH 7.5, 150 mM NaCl, 2 mM MgCl_2_, 0.5 mM TCEP, 10 μM ATP-γS, 2 μM GDC-0623, 10 μM GMP-PNP). Peak fractions containing the complex were pooled and concentrated to a volume of 500 μl and an approximate concentration of 5 μM for the protein complex. Freshly prepared bis(sulfosuccinimidyl)suberate (BS3) solution was added to the concentrated complex to a final concentration of 1 mM and incubated for 45 min at room temperature, followed by quenching of the crosslinking reaction by addition of 100 mM Tris-HCl (pH 7.5) for 15 min. Finally, the cross-linked KRAS/BRAF/MEK1/14-3-3 complex was subjected to a “polishing” round of size-exclusion chromatography in SEC buffer to remove unreacted BS3 and any aggregated or oligomerized complex.

### Cryo-EM data acquisition and processing

KRAS^GTPase^/BRAF^WT^/MEK1^SASA^/14-3-3 complex in SEC buffer was applied to glow-discharged holey carbon grids (Quantifoil R1.2/1.3, 400 mesh) and vitrified using a Leica EM GP. Frozen hydrated samples were imaged on an FEI Titan Krios at 300 kV with a Gatan Quantum Image Filter with K3 direct detection camera in super-resolution mode with a total exposure dose of ∼45 electrons. 40 frames per movie were collected at a magnification of 105,000×, corresponding to 0.85 Å per pixel. 8148 micrographs were collected at defocus values ranging from -1.8 to -2.8 μm. Crosslinked KRAS4b^1-169^/BRAF^WT^/MEK1^SASA^/14-3-3 complex with BS3 was vitrified and imaged as above on the Krios in counting mode. Total dose of 50 electrons and 50 frames per movie were collected at 0.825 Å per pixel with defocus value ranging -1.5 to -2.5 μm from two data collections of 11178 and 11144 images. Details of the data collection and dataset parameters are summarized in Extended Data Table 1. Dose-fractionated images were gain normalized, aligned, dose-weighted and summed using MotionCor2 (crosslinked sample) or the motion correction implementation within Relion (non-crosslinked sample)^29^. Contrast transfer function (CTF) and defocus value estimation were performed using CTFFIND4^30^ for the crosslinked sample and Patch CTF estimation within cryoparc for the non-crosslinked data^31^. Workflow for the single particle reconstructions from the non-crosslinked and crosslinked samples is shown schematically in Supplementary Figures 1 and 2, respectively. In brief, particle picking was carried out using crYOLO ^32^ followed by initial 2D classification within Relion^33^ to give 570,743 and 3,989,095 particles for the non-crosslinked and crosslinked samples respectively. Multiple rounds of 2D and 3D classification was then carried out on the sample to remove “junk” particles and to identify particles with distinct KRAS positions. This resulted in 69,377 particles in the non-crosslinked samples in the “Up” conformation which led to a 4.3Å reconstruction following Non-uniform refinement in cryosparc^34^. For the crosslinked sample, 190,489 particles were identified with KRAS in the “Front” position resulting in a 3.9Å reconstruction from Non-uniform refinement. Structural biology applications other than cryosparc used in this project were compiled and configured by SBGrid. Models were fit into the map using Coot^35^ and further refined with PHENIX^36^. Statistics for the final refinement are presented in Extended Data Table 1. A representative image and 2D class averages for the crosslinked sample, together with “gold-standard” Fourier shell correlation plots and heat maps showing the distribution of particle orientations for both the KRAS-up and KRAS-front structures are presented in Supplementary Figure 3.

### BRAF activity and KRAS/BRAF/MEK/ERK pathway reconstitution assays

BRAF activity assays with autoinhibited complex (BRAF^WT^/MEK^AA^/14-3-3 or BRAF^WT^/MEK^WT^/14-3-3) and active dimer (BRAF^WT^/14-3-3) were performed in assay buffer (20 mM HEPES, pH 7.4, 150 mM NaCl, 10 mM MgCl2, 1 mM TCEP, 1 mM NaVO_4_) at 25 °C. The final enzyme complex concentration in the reactions was 5 nM. Equimolar wild type MEK1 protein was added to BRAF^WT^/MEK^AA^/14-3-3 and BRAF^WT^/14-3-3 complexes for further reaction. ERK2 protein at a concentration of 2 μM was used as substrate. KRAS at a concentration of 10 nM of (a 2:1 molar ratio with the BRAF complex) was added to reaction mixture in RAS activation experiments. Reactions were initiated by adding ATP to a final concentration of 1 mM and quenched at the indicated time points by adding 2X SDS sample buffer followed by heat inactivation for 5 min at 85 °C. Assay results were analyzed by western blot with Anti-ERK (Cell signaling technology, #9102) and pERK(Cell signaling technology, #45899) antibody. For activation experiments that included liposomes, liposomes were prepared by hydrating freeze-dried Brain Phosphoserine (Avanti, #840032) with a buffer containing 20mM Hepes, pH 7.4, and 150mM NaCl to a concentration of 2.5mg/ml. The hydrated lipids were frozen in liquid N_2_ and thawed at 50°C a total of 5 times, then passed through a mini-extruder 10 times to make large unilamellar vesicles. The activation experiments with liposomes were performed in the presence of 0.1 mg/ml liposomes (final concentration), and the enzyme reactions and western blotting were carried out as described above.

### Preparation of KRAS and BRAF proteins for microscale thermophoresis studies

DNA encoding residues 1-169 of KRAS was cloned into a modified pET vector for expression with a TEV-cleavable N-terminal His_6_-tag in E. Coli. The sequence of the TEV cleavage site was modified from ENLYFQS to ENLYFQC so that after cleavage to remove the His_6_-tag, KRAS is left with an N-terminal cysteine residue for purposes of fluorescent labeling (see below). The BRAF RBD domain (residues 151-232) was also cloned into the modified pET vector for expression with a TEV-cleavable N-terminal His_6_-tag in E. Coli. The resulting plasmids were transformed into BL21(DE3) cells. Protein expressions were induced by the addition of 1 mM isopropyl-β-D-thiogalactoside at 20°C for 18 hr. Cells were harvested by centrifugation and resuspended in 20 mM HEPES (pH 7.0), 200 mM NaCl, 10 mM imidazole. After sonification, the cell lysate was applied to Ni-NTA agarose beads (Qiagen) and then eluted with 300 mM imidazole. Longer BRAF constructs that included the CRD region (residues 32-320, 39-434, and 233-766) were cloned into the modified baculovirus transfer vector pAc8 with a TEV-cleavable N-terminal His_6_-tag. These CRD-containing BRAF proteins were expressed by baculoviral infection of Sf9 insect cells and purified by Ni-NTA affinity essentially as described above. Full-length BRAF/14-3-3 and BRAF/MEK1/14-3-3 complexes in active and autoinhibited states, respectively, were prepared as previously described ^6^.

### Fluorescent labeling and GMP-PNP loading of KRAS for microscale thermophoresis studies

KRAS was labeled with Alexa Fluor 647 at its amino terminus following the protocol described in Jiang et al. ^37^ Briefly, to expose the N-terminal cysteine residue, the His_6_ tag was removed by TEV protease. During the cleavage procedure, a ‘one-pot’ reaction with MESNA (Sodium 2-mercaptoenthanesulfonate, Sigma Aldrich M1511) and Alexa Fluor™ 647 NHS Ester (Invitrogen cat# A20006) was initiated. This procedure yielded KRAS selectively labeled with Alex Fluor 647 via an amide bond with the N-terminal cysteine residue. After the labeling reaction, the labeled KRAS protein was purified by size-exclusion chromatography using 16/60 Superdex75 increase column (GE Healthcare) pre-equilibrated with 20 mM HEPES (pH 7.0), 150 mM NaCl, 1 mM TCEP.

As purified, KRAS is bound primarily to GDP. In order to load with the non-hydrolyzable GTP analog GMP-PNP for binding studies, we carried out an exchange procedure as described previously. ^38^ Briefly, purified KRAS protein was mixed with GMP-PNP (molar ratio of 10:1 GMP-PNP:KRAS) and calf intestinal alkaline phosphatase (NEB cat# M0290, 3 units per mg of KRAS). The reaction mixture was incubated for 3 hours at room temperature and then purified by size-exclusion chromatography on a 10/300 Superdex 75 GL column (GE Healthcare) pre-equilibrated with 100 mM HEPES (pH 7.0), 150 mM NaCl, 1 mM Tris(2-carboxyethyl)phosphine hydrochloride (TCEP). GMP-PNP loaded KRAS was then concentrated, flash frozen, and stored at -80°C until used.

### Microscale thermophoresis measurements

Binding affinities of the Alexa Fluor™ 647 labeled GMPPNP bound KRAS and BRAF proteins were measured using microscale thermophoresis (MST). Before the MST experiment, A Tween 20 detergent was added into all samples to 0.05%. 10 μL of serially diluted BRAF proteins from 10 pM to 1 μM were loaded into 8 PCR tubes. Then, 10 μL of 5 nM N-terminal labeled KRAS mixed into each reaction tubes. The binding affinity measurements were carried out using the modified manufacturer’s protocol (Monolith NT.115pico, NanoTemper Technologies) in the Center for Macromolecular Interactions in Harvard Medical School (Boston, MA).

### Crosslinking mass spectrometry studies

Complexes crosslinked with BS3 were denatured in 8 M urea and subjected to reduction and carbamidomethylation with TCEP and iodoacetamide. Samples were diluted to 2 M urea and digested with trypsin (1:50 ratio of trypsin:protein) overnight at 32°C. Digestions were brought to 1% formic acid (FA) and peptides were dried by vacuum centrifugation and desalted over a C18 column. Digests were resuspended in 5% acetonitrile (MeCN)/1% FA and analyzed on an Ultimate 3000 RSLCnano system coupled to an Orbitrap Eclipse mass spectrometer (Thermo Scientific). Peptides were separated across a 55-min linear gradient of 7-22% MeCN in 1% FA, followed by 15 min to 45% MeCN over a 50-cm C18 column (ES803A, Thermo Scientific, Waltham, MA, USA) and electrosprayed (2.15 kV, 300°C) into the mass spectrometer with an EasySpray ion source (Thermo Scientific). Precursor ion scans (300-2,000 *m/z*) were obtained in the orbitrap at 120,000 resolution in profile (RF lens % = 30, Max IT = 100 ms, 1 microscan). Fragment ion scans were acquired in the orbitrap at 30,000 resolution (1.6 *m/z* isolation window, HCD at 30% NCE, 5e^4^ AGC).

Raw data were searched against KRAS, BRAF and MEK1 construct sequences, as well as endogenous Sf9 14-3-3ε and 14-3-3ζ sequences for BS3 crosslinked peptides using Protein Prospector Batch-Tag (https://prospector.ucsf.edu/) ^39,40^ permitting mass accuracy of ±5 ppm for precursors and ±10 ppm for fragment ions, two missed cleavages by trypsin, oxidation of Met, carbamidomethylation of Cys, phosphorylation of Ser/Thr/Tyr, and BS3 monolinked Lys. Crosslinked hits were filtered for spectra with Protein Prospector score differences >15. Resulting spectra were manually inspected to ensure the presence of abundant fragment ions from both crosslinked peptides. A filtered peptide spectral match (PSM) file and representative spectra of manually confirmed inter- and intra-molecular crosslinks are presented in Supplementary Data Files 1 and 2, respectively, in the Supplemental Information.

---

**Extended Data Figure 1.**
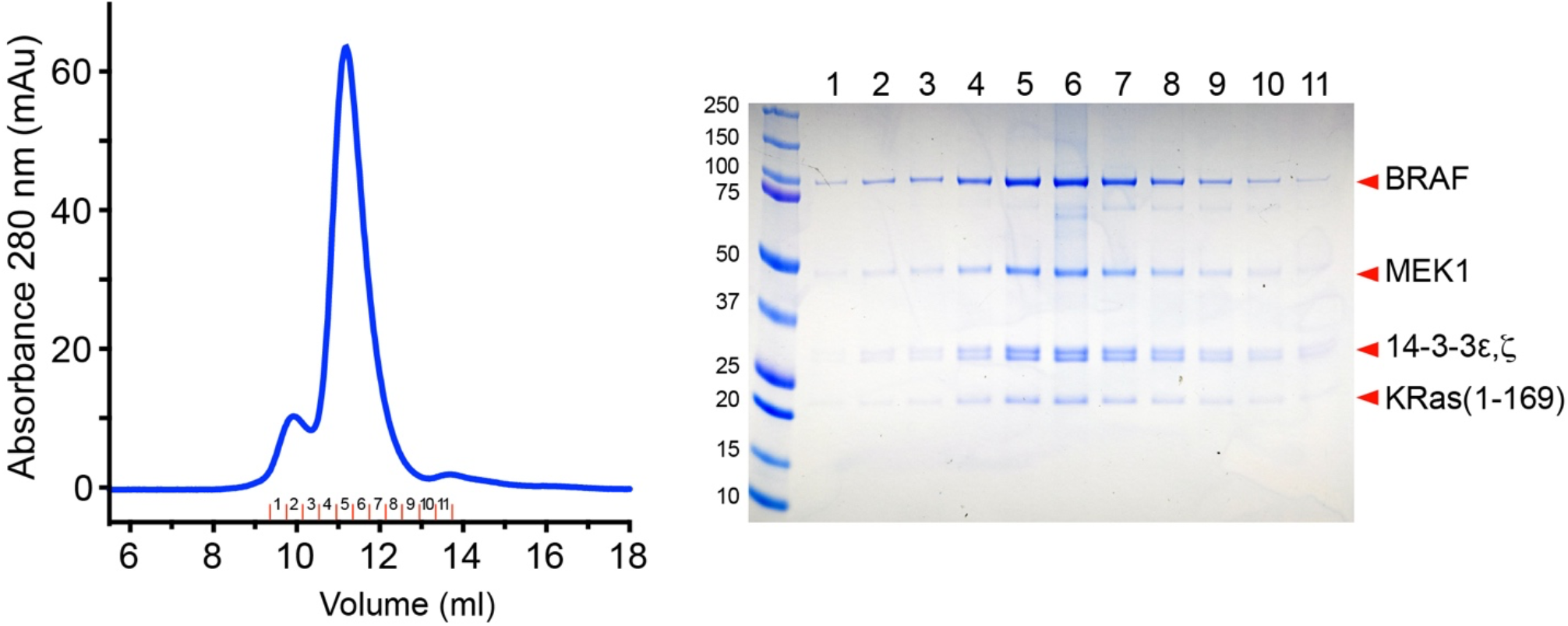
Size-exclusion chromatography of the KRAS/BRAF/MEK1/14-3-3 complex. Elution profile of the complex on Superdex S200 is shown on the left, and a Coomassie-stained SDS-PAGE gel of the indicated fractions is shown on the right.

**Extended Data Figure 2.**
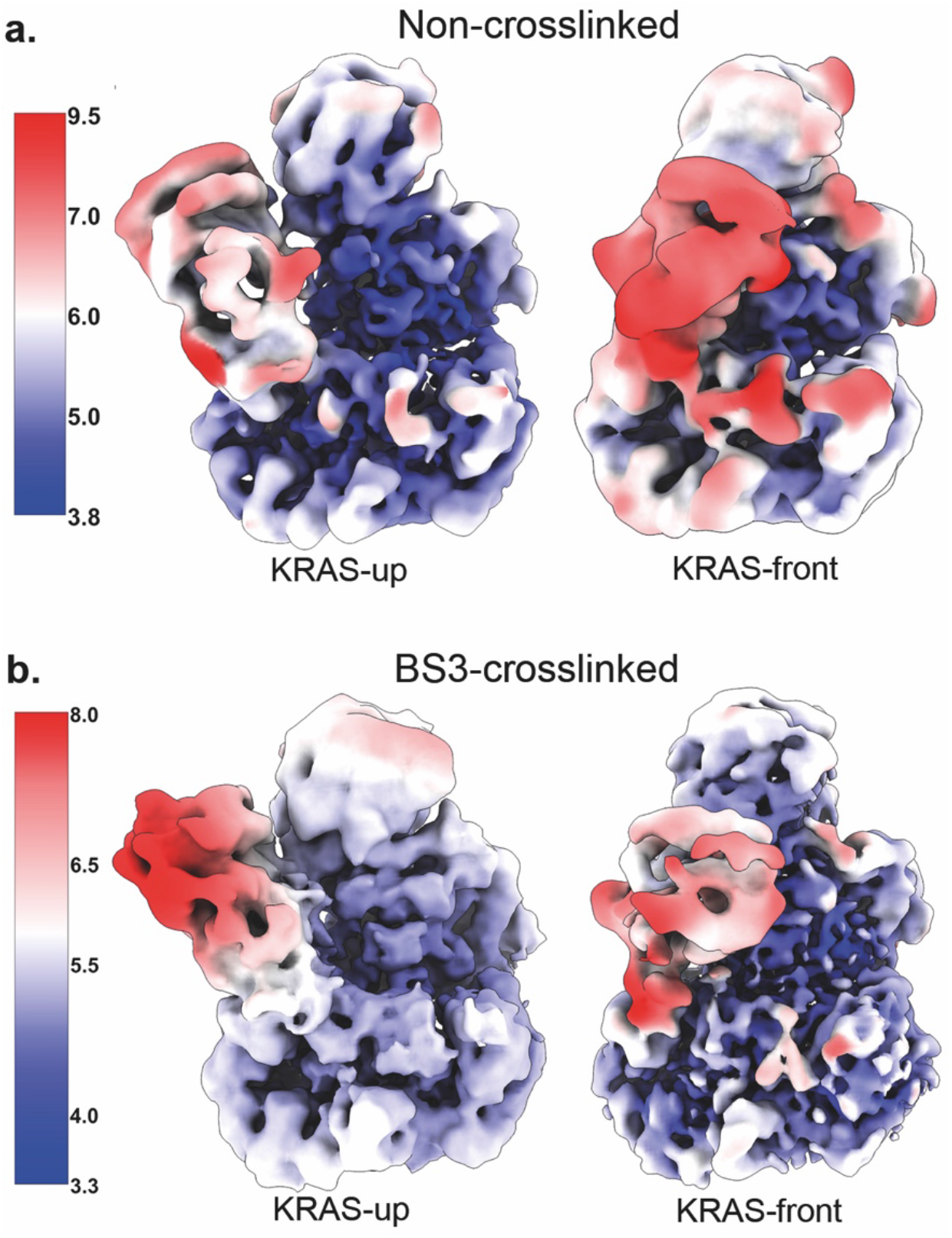
Cryo-EM density maps obtained from single particle reconstructions from crosslinked and non-crosslinked KRAS/BRAF/MEK1/14-3-3 samples. 3D classification of particles allowed identification of two conformations of the complex, KRAS-up and KRAS-front, in both non-crosslinked (a) and BS3-crosslinked (b) samples. Maps are colored by resolution (Å) as indicated by the bars on the left.

**Extended Data Figure 3.**
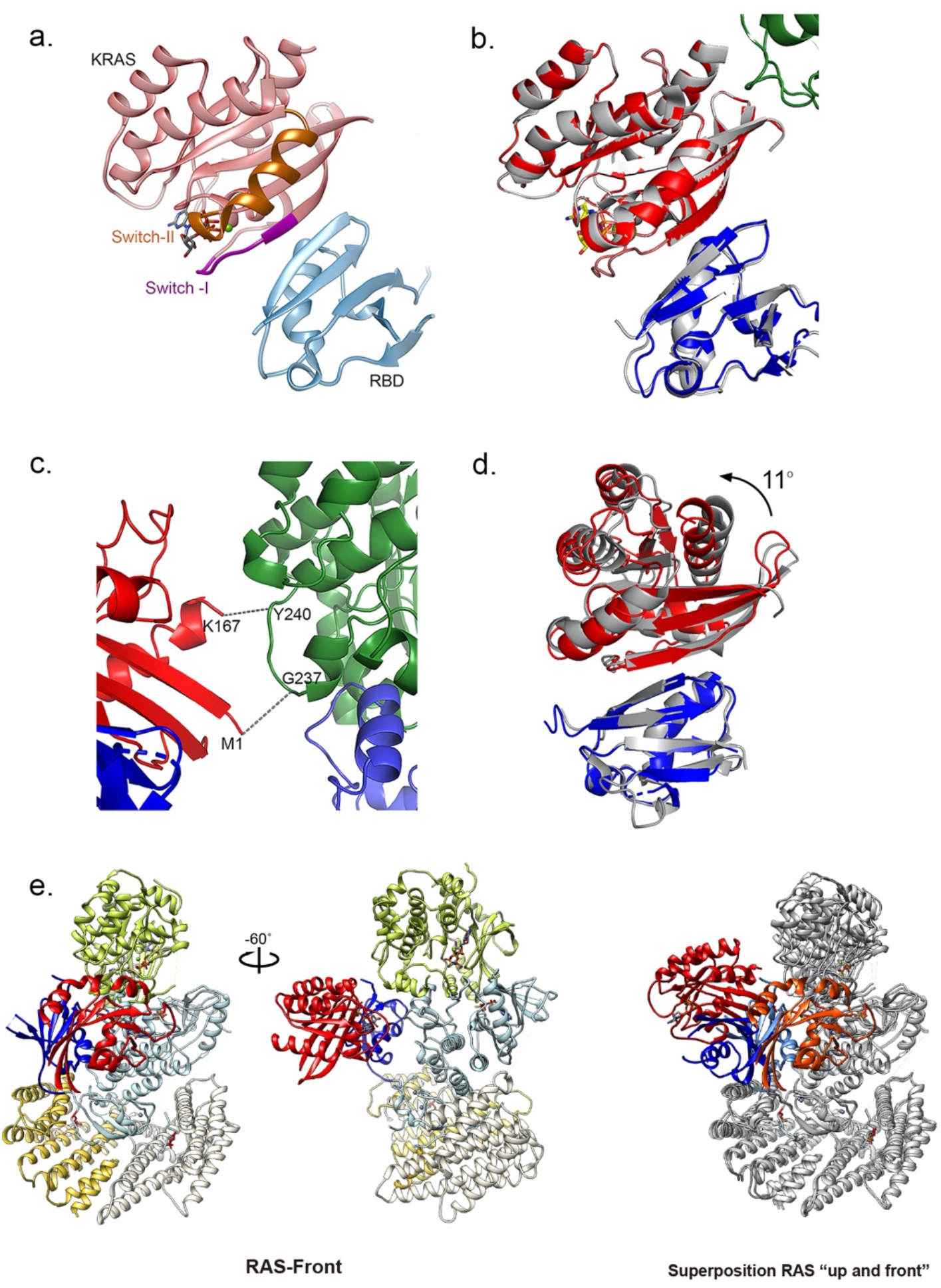
Additional views and comparisons of the KRAS/RBD module. a, Ribbon diagram showing the interaction between KRAS and the BRAF RBD domain in the KRAS-up structure. The Switch-I and Switch-II regions of KRAS are highlighted in purple and orange, respectively. b, Superposition of the KRAS and RBD region of the KRAS-up structure with the previously determined structure of KRAS bound to the isolated RBD domain of CRAF (gray ribbon, PDB entry 6VJJ). c, Region of contact between KRAS (red) and MEK1 (green) in the KRAS-up structure. d, Superposition of the KRAS and RBD region of the KRAS-front structure with the previously determined structure of KRAS bound to the isolated RBD domain of CRAF (color-coded as in panel b). Superposition based on the RBD domains reveals a change in the relative orientation of KRAS by approximately 11°. e, Two views of the KRAS-front structure show the contact of the KRAS/RBD module at the interface between the BRAF and MEK1 kinase domains. Superposition of the KRAS-up and -front structures (right panel) reveals a ∼165° difference in the relative orientation of the KRAS/RBD module.

**Extended Data Figure 4.**
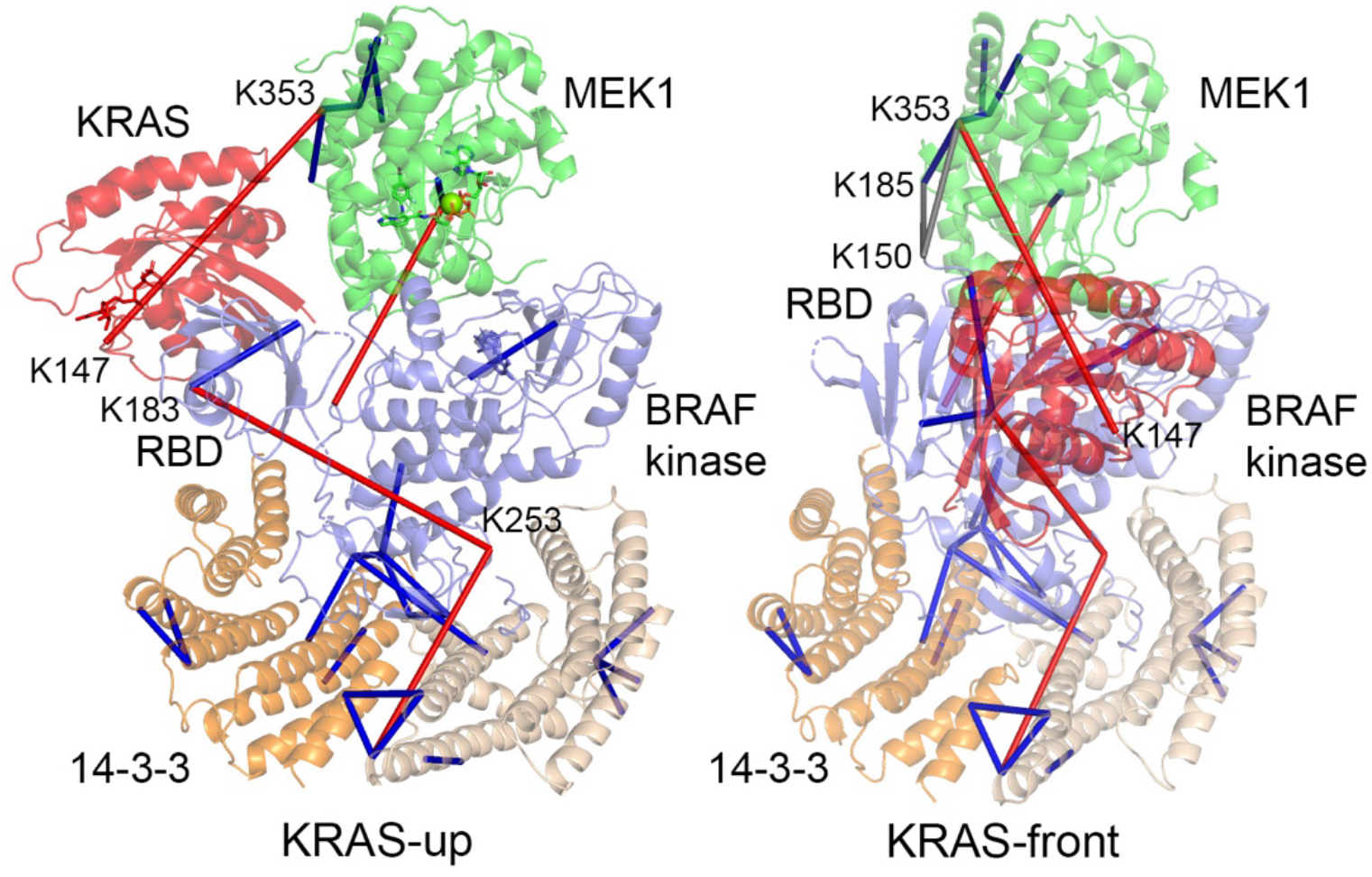
Crosslinking mass spectrometry analysis of the BS3-crosslinked KRAS/BRAF/MEK1/14-3-3 sample. Crosslinks identified by mass spectrometry analysis of the BS3-treated sample are mapped onto the KRAS-up (left) and KRAS-front (right) structures. Residue numbers are given for inter-domain crosslinks to KRAS and the RBD domain. Crosslinks that are consistent with the structures (Cα to Cα distance of 20Å or less) are shown in dark blue, those that exceed this cutoff are shown in red. Two crosslinks between BRAF residue K150, which is just N-terminal to the RBD domain, and MEK1 residues K185 and K353 are indicated in gray and are consistent with the KRAS-front conformation.

**Extended Data Figure 5.**
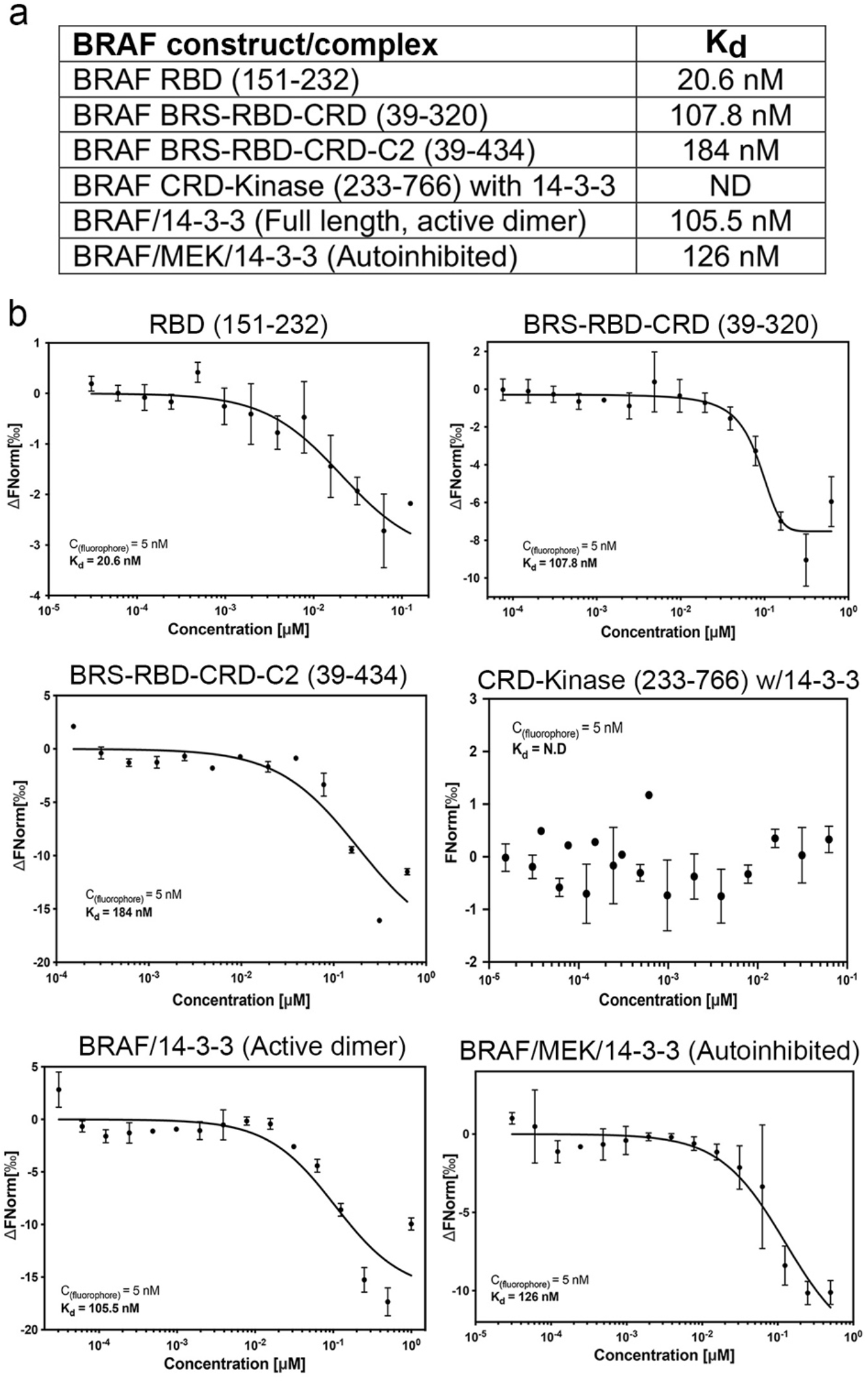
Microscale thermophoresis studies of binding of KRAS to BRAF fragments and complexes. a, Measured affinity (dissociation constant, K_d_) of GMP-PNP loaded KRAS^GTPase^ for the indicated BRAF fragments or for full-length BRAF bound to 14-3-3 in the active, dimeric state or full-length BRAF/MEK1/14-3-3 in the autoinhibited state. b, Microscale thermophoresis titration curves used to determine the dissociation constants in panel a.

