## Supplemental Figures for "Cryo-EM structure of a RAS/RAF recruitment complex"

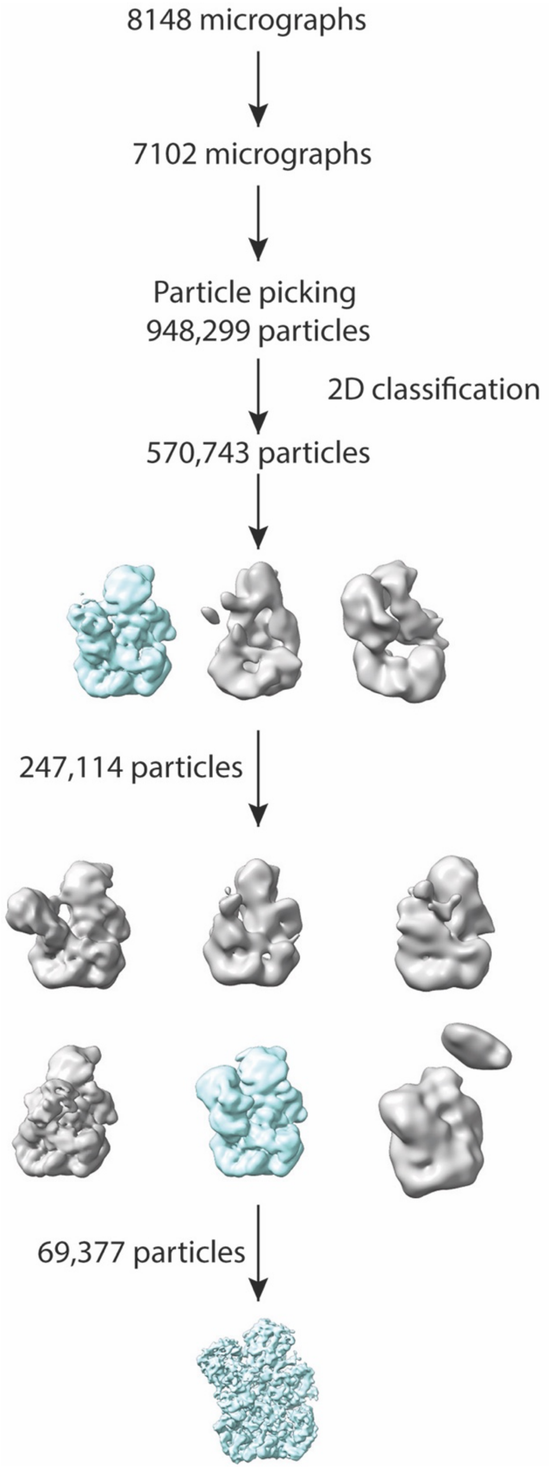


**Supplementary Figure 1. Flowchart for single particle reconstruction of the KRAS/BRAF/MEK1/14-3-3 structure in the "KRAS-up" conformation (non-crosslinked).**

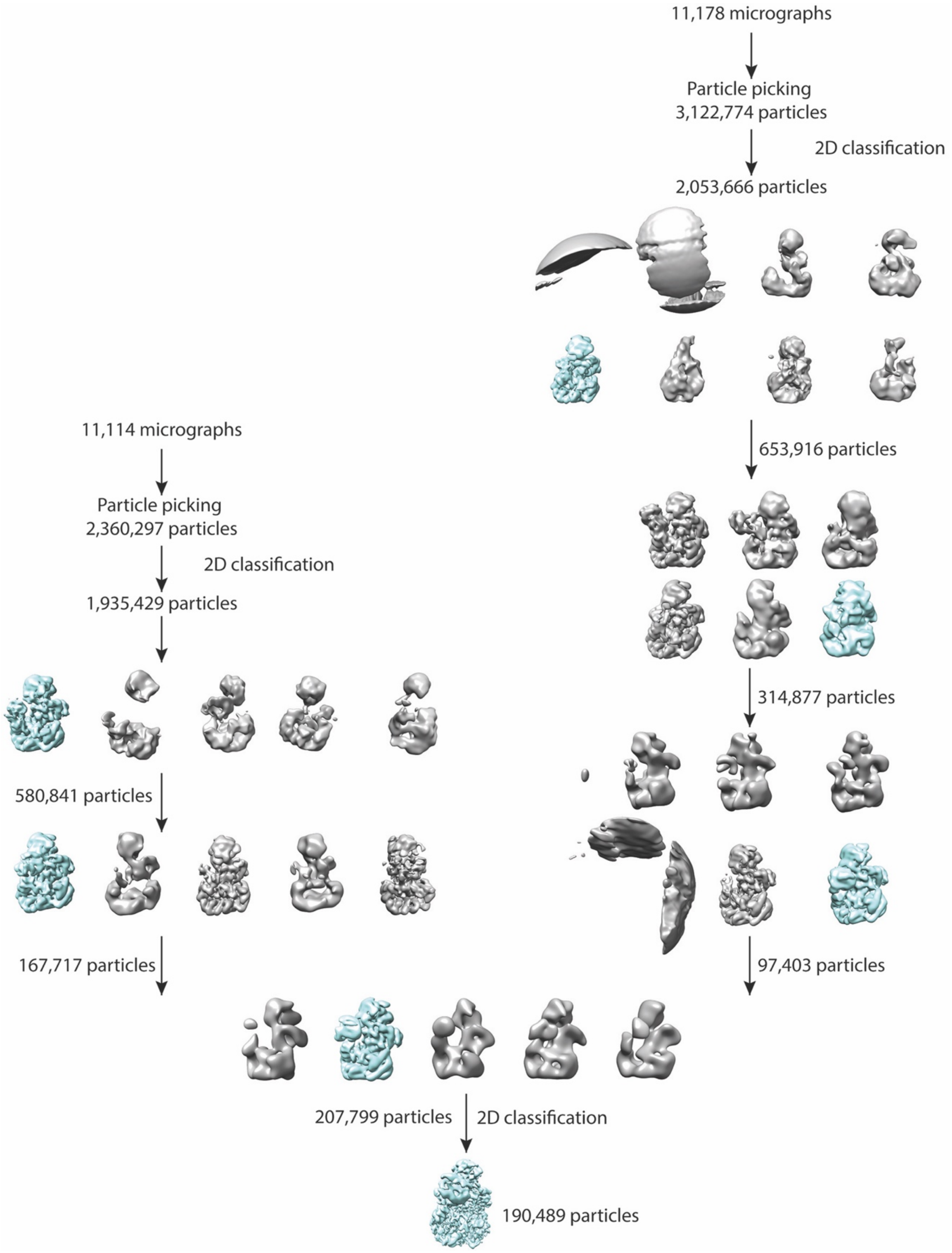


**Supplementary Figure 2. Data processing flowchart the KRAS/BRAF/MEK1/14-3-3 structure in the "KRAS-front" conformation (BS3-crosslinked).** Particle stacks were merged from data sets collected from two grids prepared from the same protein sample.

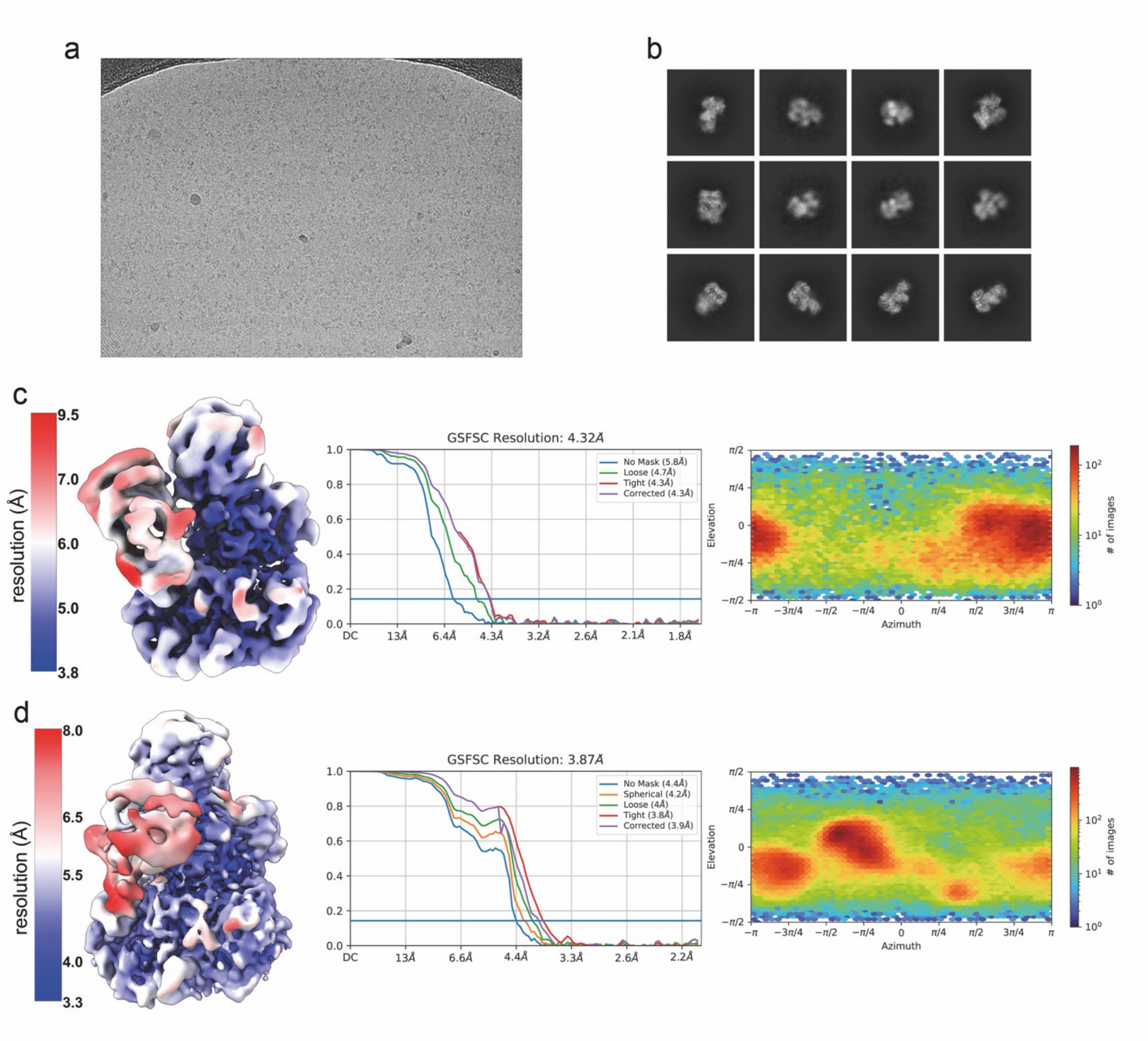


**Supplementary Figure 3. Supporting information for cryo-EM structure determination.** a, Section of a representative micrograph of the BS3-crosslinked sample. b, Representative 2D-class averages from the BS3-crosslinked sample. c and d, Cryo-EM density maps colored by resolution for the KRAS-up and KRAS-front structures, respectively, together with corresponding gold-standard Fourier shell correlation curves and heatmaps showing distribution of particle orientations.
