## Supplemental Data 2 for "Cryo-EM structure of a RAS/RAF recruitment complex"

MS/MS spectra of **A**) BRAF–14-3-3ε, **B**) BRAF–14-3-3ζ, **C**) BRAF–MEK, **D**) KRAS–MEK, **E**) 14-3-3ε–14-3-3ζ, F) 14-3-3ε–14-3-3ε, **G**) MEK–MEK, **H**) 14-3-3ζ–14-3-3ζ, and **I**) BRAF– BRAF crosslinked peptides. B and y ions originating from each crosslinked peptide are colored in blue or green, corresponding to the peptide sequence of the same color. Cases in which fragmentation occurred across the linker are indicated as “P+L” or “P+LK”, ions consisting of the peptide and linker or the peptide, linker and linked Lys residue of the opposing peptide, respectively.

**A**

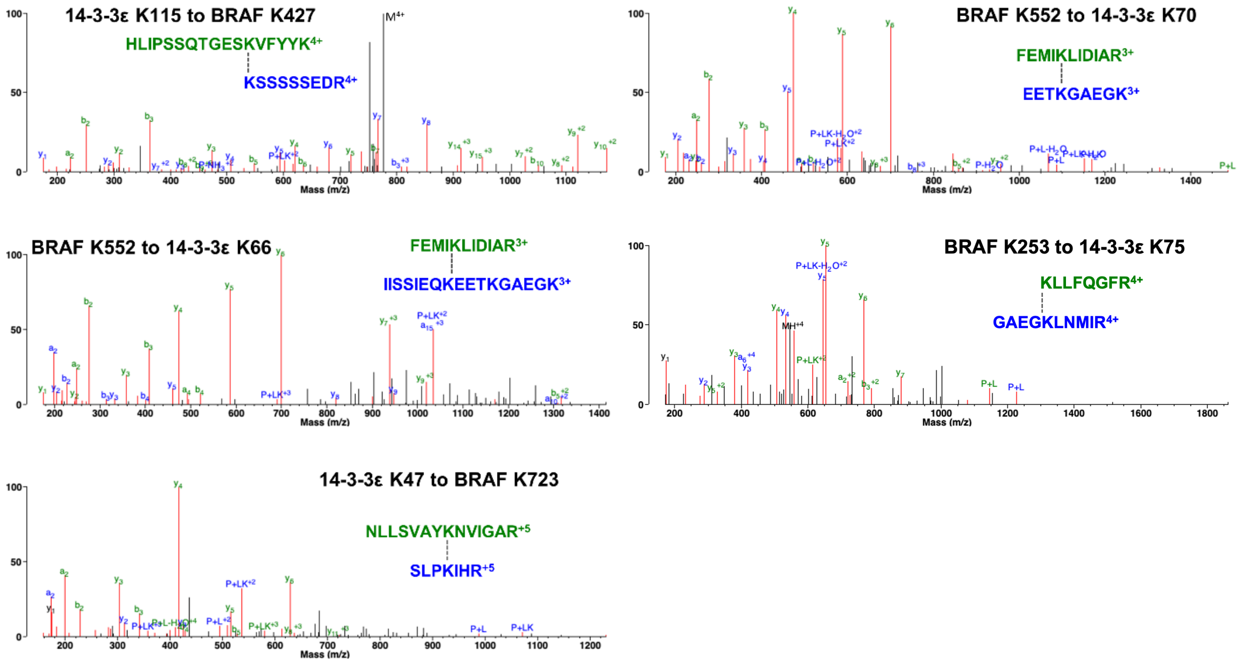


**B**

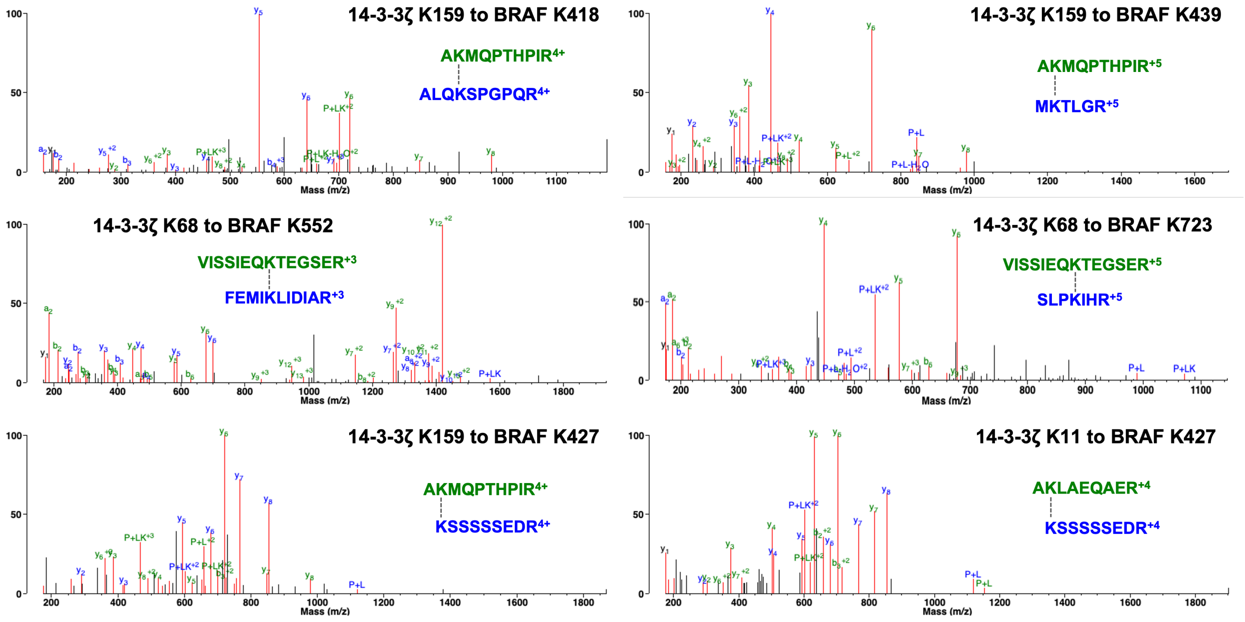

C

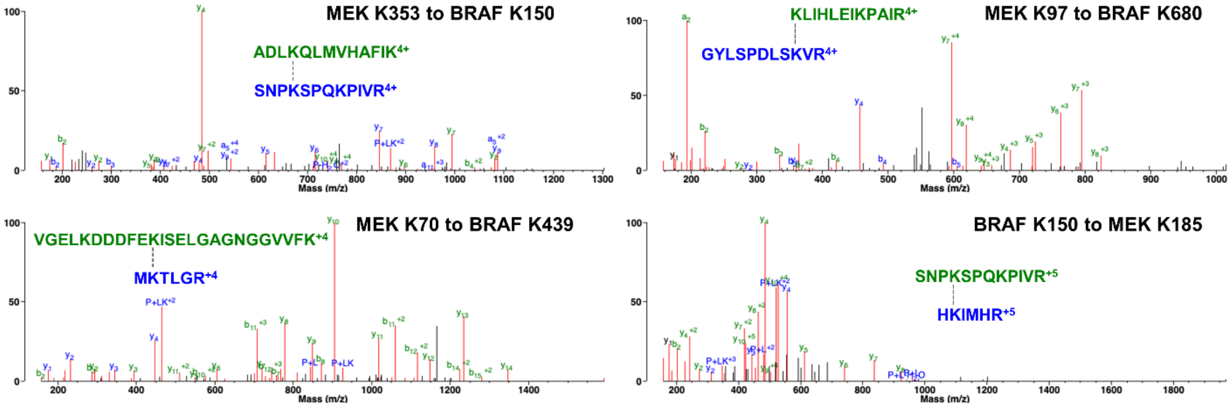
MEK K353 to BRAF K150
ADLKQLMVHAFIK4+
SNPKSPQKPIVR4+
MEK K97 to BRAF K680
KLIHLEIKPAIR4+
GYLSPDLSKVR4+
MEK K70 to BRAF K439
VGELKDDDFEKISELGAGNGGVVFK+4
MKTLGR+4
BRAF K150 to MEK K185
SNPKSPQKPIVR+5
HKIMHR+5
Mass (m/z)

D

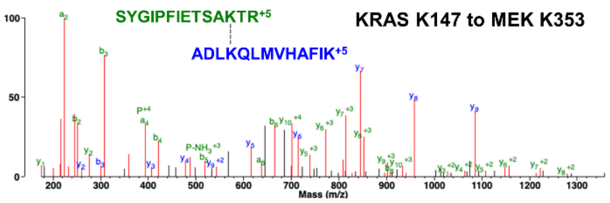
KRAS K147 to MEK K353
SYGIPFIETSAKTR+5
ADLKQLMVHAFIK+5
Mass (m/z)

E

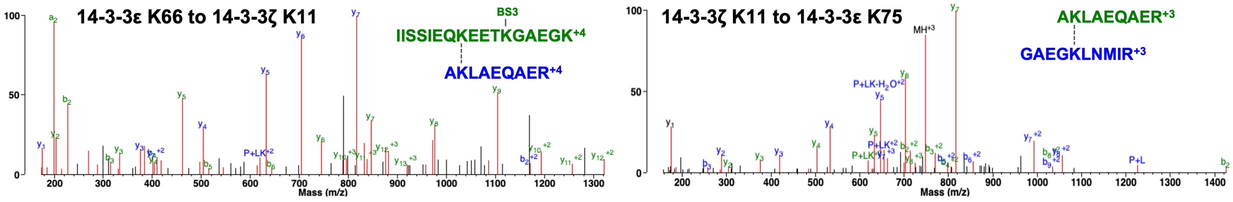
14-3-3ε K66 to 14-3-3ζ K11
BS3
IISSIEQKEETKGAEGK+4
AKLAEQAER+4
14-3-3ζ K11 to 14-3-3ε K75
AKLAEQAER+3
GAEGKLNMIR+3
Mass (m/z)

F

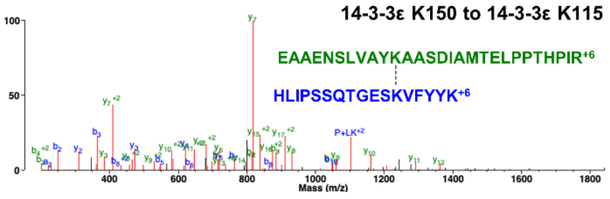
14-3-3ε K150 to 14-3-3ε K115
EAAENSLVAYKAASDIAMTELPPTHPIR+6
HLIPSSQTGESKVFYYK+6
Mass (m/z)

G

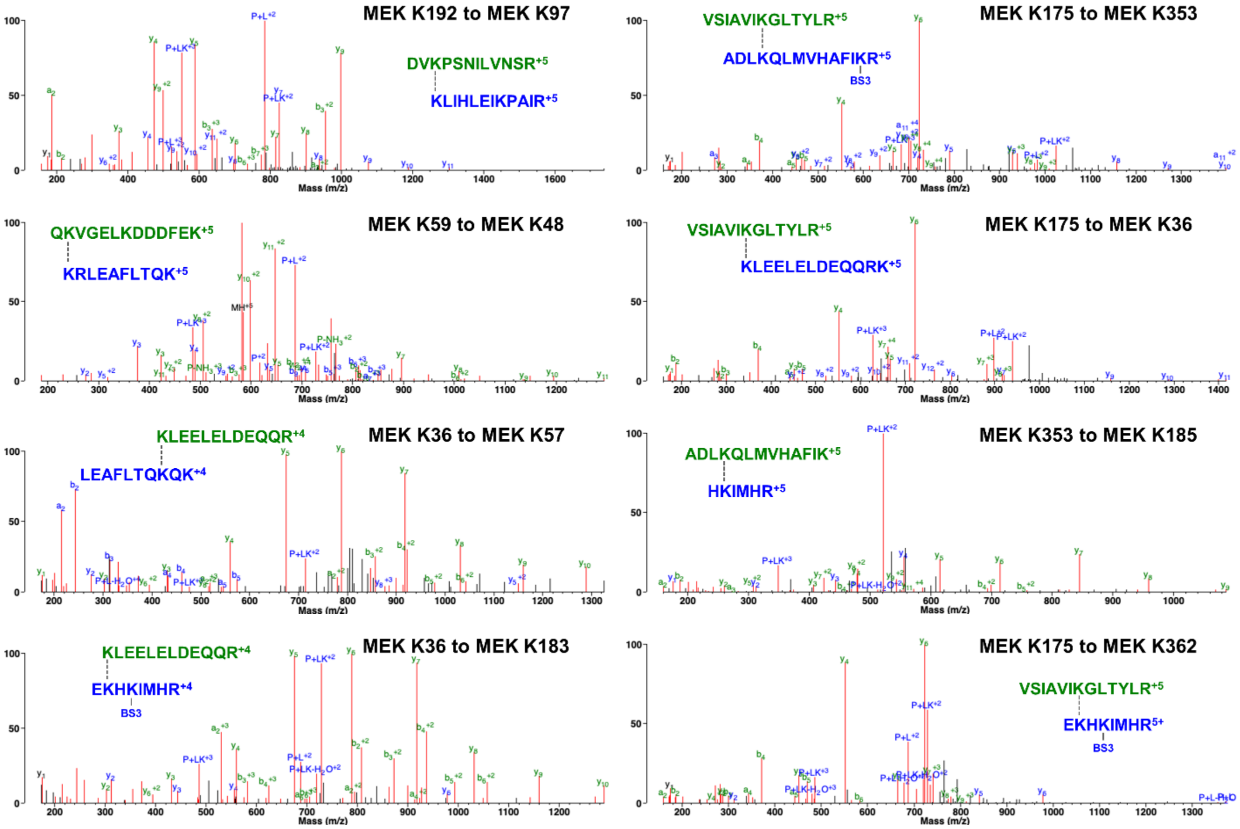
MEK K192 to MEK K97
DVKPSNILVNSR+5
KLIHLEIKPAIR+5
MEK K175 to MEK K353
VSIAVIKGLTYLR+5
ADLKQLMVHAFIKR+5
BS3
MEK K59 to MEK K48
QKVGELKDDDFEK+5
KRLEAFLTQK+5
MEK K175 to MEK K36
VSIAVIKGLTYLR+5
KLEELELDEQQRK+5
MEK K36 to MEK K57
KLEELELDEQQR+4
LEAFLTQKQK+4
MEK K353 to MEK K185
ADLKQLMVHAFIK+5
HKIMHR+5
MEK K36 to MEK K183
KLEELELDEQQR+4
EKHKIMHR+4
BS3
MEK K175 to MEK K362
VSIAVIKGLTYLR+5
EKHKIMHR5+
BS3
Mass (m/z)

H

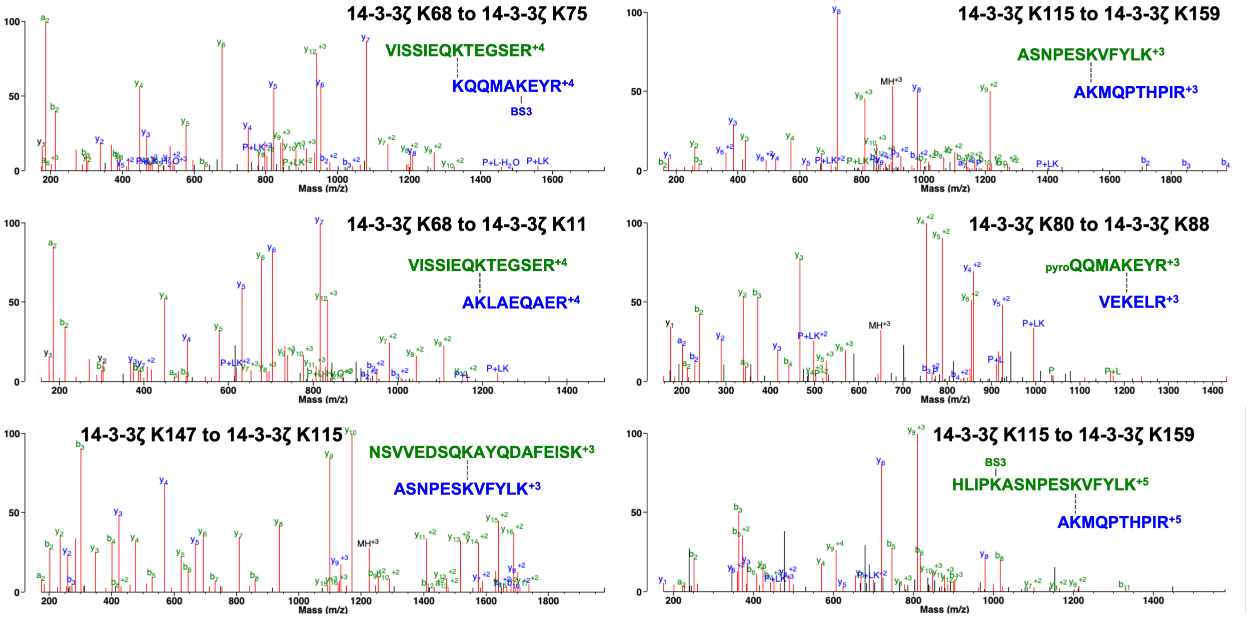
14-3-3ζ K68 to 14-3-3ζ K75
VISSIEQKTEGSER+4
KQQMAKEYR+4
BS3
14-3-3ζ K115 to 14-3-3ζ K159
ASNPESKVFYLK+3
AKMQPTHPIR+3
14-3-3ζ K68 to 14-3-3ζ K11
VISSIEQKTEGSER+4
AKLAEQAER+4
14-3-3ζ K80 to 14-3-3ζ K88
pyroQQMAKEYR+3
VEKELR+3
14-3-3ζ K147 to 14-3-3ζ K115
NSVVEDSQKAYQDAFEISK+3
ASNPESKVFYLK+3
14-3-3ζ K115 to 14-3-3ζ K159
BS3
HLIPKASNPESKVFYLK+5
AKMQPTHPIR+5
Mass (m/z)

I

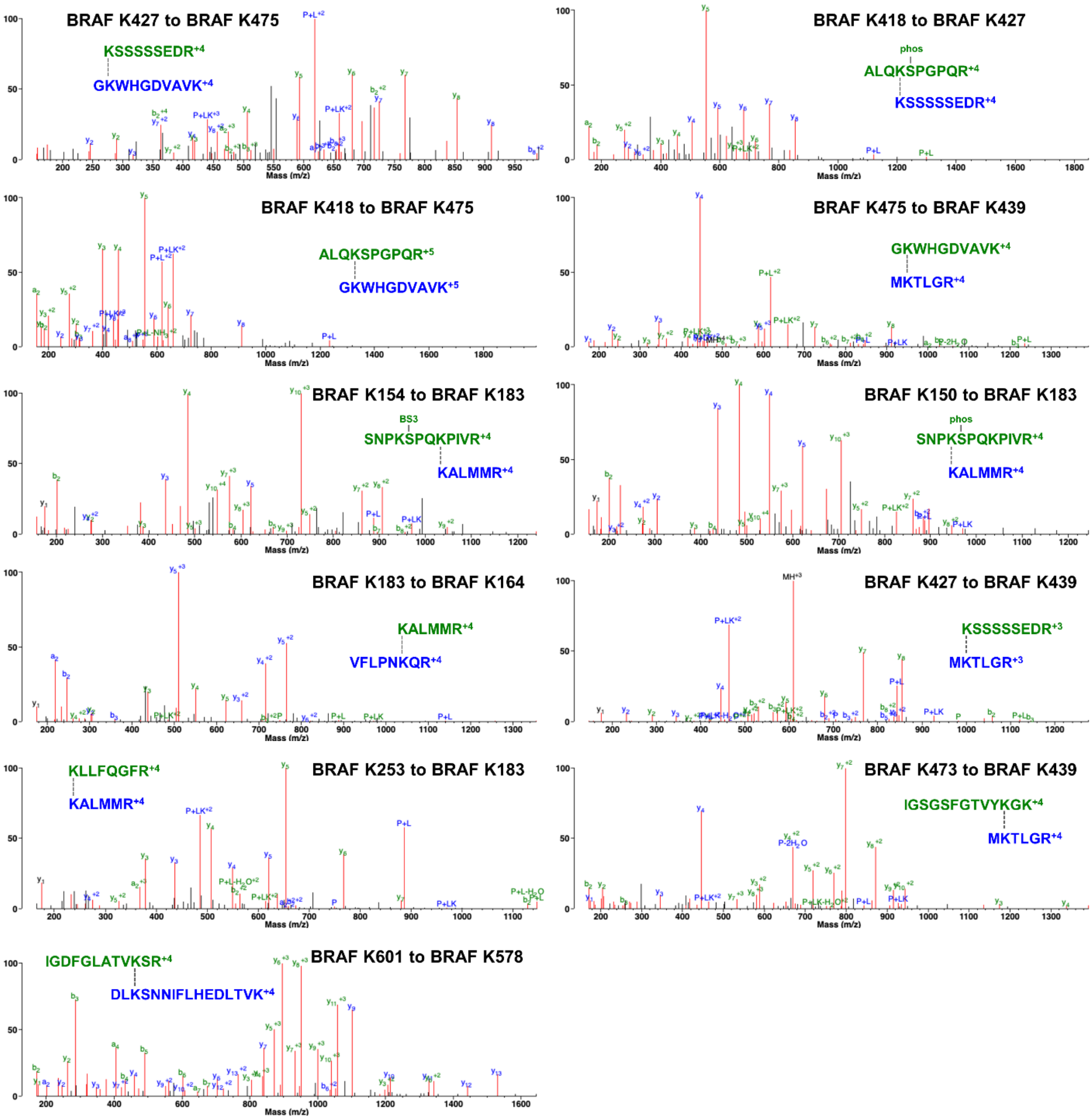
BRAF K427 to BRAF K475
KSSSSSEDR+4
GKWHGDVAVK+4
BRAF K418 to BRAF K427
phos
ALQKSPGPQR+4
KSSSSSEDR+4
BRAF K418 to BRAF K475
ALQKSPGPQR+5
GKWHGDVAVK+5
BRAF K475 to BRAF K439
GKWHGDVAVK+4
MKTLGR+4
BRAF K154 to BRAF K183
BS3
SNPKSPQKPIVR+4
KALMMR+4
BRAF K150 to BRAF K183
phos
SNPKSPQKPIVR+4
KALMMR+4
BRAF K183 to BRAF K164
KALMMR+4
VFLPNKQR+4
BRAF K427 to BRAF K439
KSSSSSEDR+3
MKTLGR+3
KLLFQGFR+4
KALMMR+4
BRAF K253 to BRAF K183
BRAF K473 to BRAF K439
IGSGSFGTVYKGK+4
MKTLGR+4
IGDFGLATVKSR+4
DLKSNNIFLHEDLTVK+4
BRAF K601 to BRAF K578
Mass (m/z)
